# Distinct effects of sensory and genetic risk factors for psychosis on auditory cortical temporal acuity in a mouse model of 22q11.2 Deletion Syndrome

**DOI:** 10.64898/2026.03.26.714426

**Authors:** Chen Lu, Jennifer F. Linden

## Abstract

**Background:** Peripheral hearing loss is associated with auditory hallucinations and increased risk for psychotic disorder, particularly in genetically vulnerable individuals. Moreover, both hearing loss and schizophrenia disrupt auditory temporal acuity, a sensitive measure of auditory brain function. Here, we used mouse models of hearing loss and schizophrenia to reveal how neural mechanisms of auditory cortical temporal acuity depend on sensory and genetic risk factors for psychosis.

**Methods:** We quantified auditory cortical temporal acuity in mice with or without hearing loss and with or without the 22q11.2 deletion — one of the strongest known genetic risk factors for schizophrenia. Cortical single-unit and population activity were recorded in awake mice (N = 23) during presentations of loudness-adjusted gap-in-noise stimuli with varying durations of silent gap, which are commonly used to assess auditory temporal acuity in humans.

**Results:** Both hearing loss and the 22q11.2 deletion disrupted auditory cortical temporal acuity, but through distinct mechanisms. Hearing loss broadly degraded temporal acuity at the level of single-unit responses and neural population activity. In contrast, the 22q11.2 deletion selectively impaired gap duration thresholds in regular-spiking (putative excitatory) but not fast-spiking (putative inhibitory) neurons. Mice with comorbid hearing loss and 22q11.2 deletion exhibited both abnormalities in auditory cortical temporal acuity.

**Conclusions:** Sensory and genetic risk factors for psychosis can disrupt auditory cortical temporal acuity via distinct mechanisms that remain partially dissociable even under comorbid conditions. These findings underscore the importance of accounting for hearing loss comorbidity when interpreting auditory cortical dysfunction in psychotic disorder.

## Introduction

Clinical studies have shown that psychosis impairs auditory processing, including frequency discrimination, spatial localization, deviant detection, and prepulse inhibition (1). In addition, patients with psychosis exhibit reduced hearing sensitivity and difficulty hearing in noisy environments (2,3). Although hearing loss is recognized as a risk factor for psychosis (4), it remains unclear whether it merely co-occurs with other psychosis risk factors or independently exacerbates the risk or severity of psychotic disorder.

Temporal processing is essential for normal auditory function, supporting sound perception and speech recognition (5–7). It can be quantified using the gap-in-noise paradigm, which measures sensitivity to brief silent gaps embedded within white noise or pure tones. Patients with psychosis exhibit prolonged gap detection thresholds, indicating impaired auditory temporal acuity (8,9). In the general population, peripheral hearing loss also increases gap detection thresholds, even when sound levels are adjusted to compensate for elevated hearing thresholds (10). Here, we investigated how hearing loss and genetic risk for psychosis influence auditory cortical activity evoked by gap-in-noise stimuli, aiming to disentangle their respective contributions to auditory dysfunction.

We used *Df1/+* mice, a model of 22q11.2 deletion syndrome (22q11.2DS), to study how hearing loss and genetic risk shape auditory cortical temporal processing. The 22q11.2 deletion is one of the most common chromosomal microdeletions, with a prevalence of 1 in every 2,200 live births (11). Nearly 25% of carriers develop schizophrenia in their lifetime — compared to 0.5% in the general population — and their symptoms are similar to idiopathic schizophrenia (12). The deletion also confers a high incidence of conductive hearing loss, with over half of carriers experiencing early-onset middle-ear inflammation (13). *Df1/+* mice, which harbor a homologous chromosomal deletion, exhibit similar comorbid hearing loss in approximately 50-60% of animals and show electrophysiological brain abnormalities paralleling those observed in human deletion carriers (14,15).

In this study, we investigated how sensory and genetic risk factors for psychosis influence auditory function by comparing *Df1/+* mice with normal hearing or hearing loss to wild-type (WT) mice with normal hearing or hearing loss. We asked how auditory cortical sensitivity to brief gaps differed between animal groups with well-controlled genetic background and well-characterised hearing sensitivities. Intracranial recordings from auditory cortical neurons were obtained using Neuropixels probes while awake mice listened passively to gap-in-noise stimuli with sound levels adjusted to compensate for the hearing threshold of each mouse. Novel, robust methods for quantifying gap-duration thresholds at both the single-neuron and neuronal population levels provided rigorous metrics of auditory cortical temporal acuity in each animal group. These methods revealed that both peripheral hearing loss and the 22q11.2 deletion disrupt auditory cortical temporal acuity; however, the effects of these two risk factors differ across cortical cell types. Our findings provide a foundation for translational studies of the impact of comorbid hearing loss on psychosis pathology in humans.

## Methods

Experimental design and analysis of onset and offset responses, single-unit gap-duration thresholds, and population-level gap-duration thresholds are described briefly below. See Supplementary Information for additional details on animals, surgical procedures, auditory brain response (ABR) recordings, cortical recordings using Neuropixels probes, data processing and statistical analysis.

### Experimental design

In brief, primary auditory cortical units were recorded in awake head-fixed mice using Neuropixels 1.0 probes, while the animals listened passively to presentations of gap-in-noise stimuli with varying gap durations. The noise level was adjusted to be 20 dB above the better-ear hearing threshold for each mouse, as determined from click-evoked ABR measurements in each ear. Gap-in-noise stimuli were composed of a 250 ms noise burst followed by a silent gap of varying duration (0, 1, 2, 4, 8, 16, 32, 64, 128, or 256 ms) followed by a 100 ms noise burst. The noise was synthesized by summing discrete pre-calibrated pure tones from 8 to 64 kHz, spaced at 8 steps per octave. To avoid blurring broadband transients, no ramping was used. The inter-trial interval was 1000 ms, and there were 45 repeated trials of each of the 10 gap conditions, presented in pseudo-randomized order.

### Onset and offset response detection

The onset and offset responses of single units were detected by comparing neural responses before and after transient events in noise. Significant changes in firing rates were defined as 2 consecutive rising bins within the post-transient event period that exceeded 3 standard deviations above the mean of the pre-transient event period. For sound onset responses, we also allowed for the possibility of drops in firing rate due to sound-evoked inhibition; however, we required offset responses to manifest as an increase in firing rate because a decrease in firing rate following sound offset cannot easily be distinguished from recovery from adaptation in the absence of sound-evoked activity (16). The pre-onset period is the 100 ms before the first noise onset, and the post-onset period is 100 ms after the first noise onset. The pre-offset period is 50 ms before the second noise offset, and the post-offset period is 10 to 110 ms after the second noise offset. Onset and offset analyses were performed for all trials. A unit was classified as onset- or offset-responsive only if at least 7 out of the 10 gap conditions exhibited valid responses as defined above.

### Identification of gap duration thresholds in single units

Gap detection in single units was analyzed by comparing deviations in neural responses surrounding gaps to sustained responses to noise. The peri-gap analysis window spanned from the start of the gap to 100 ms after the gap ended, covering the entire gap and second noise period. The sustained response to noise was measured as the neural response during the last 100 ms of noise in the trial with a 0-ms gap. The root mean squared deviation (RMSD) of neural responses within the peri-gap analysis window, relative to the sustained responses, was calculated as below:

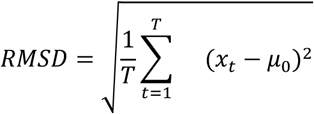

where 𝑥*_t_* is spike rate of 5-ms bin at time 𝑡, 𝜇_0_ is the mean firing rate in the last 100 ms of noise in the no-gap condition, and 𝑇 is the total number of time bins within the peri-gap analysis window.

A sigmoid function was fitted to quantify the relationship between the normalized RMSD and the binary logarithm of gap duration (in ms). In some cases, RMSD values decreased at longer gap durations or were close to zero for shorter gaps, which could reduce the quality of the sigmoid fit. In such instances, the fitting was performed again after excluding these problematic data points. The better-fitting result was used to estimate gap duration thresholds.

Two times the standard deviation of the sustained responses was defined as the threshold for significant differences. The gap duration at which the sigmoid function returned this value was defined as the gap-duration threshold of the unit.

### Identification of gap duration thresholds in neuronal populations

For population analysis, principal components analysis was applied to reduce the dimensionality of the neuronal population data. Responses from -100 ms to 900 ms after the onset of the first noise were averaged across trials with the same gap durations. The averaged responses were then concatenated for analysis. The first three principal components were used to reconstruct the trajectory of the neuronal population response.

To compare gap duration threshold differences between groups at the neuronal population level, both the distance and the area of the population trajectory representing the response to the first and second noise were calculated. Principal components analysis was performed on 50-unit subgroups that were randomly selected from the original unit pool to ensure that differences in the total number of units per group did not bias population trajectory measures. The trajectory distance was calculated by accumulating the Euclidean distances between consecutive time points. The area of the trajectory was estimated as the accumulated area formed between each pair of time points. This analysis was repeated 100 times for between-group comparisons, with different random selections of 50 units from each group.

For analysis of gap duration sensitivity, the population trajectory distance (or area) for the post-gap noise burst was divided by the trajectory distance (or area) for the pre-gap noise burst. This ratio measure provided a measure of gap duration sensitivity at the neuronal population level that was robust to differences between groups in neuronal responsiveness to noise. This approach, together with the use of sound stimuli adjusted in loudness to compensate for hearing loss on an animal-by-animal basis, ensured that differences in hearing thresholds between animal groups could not account for observed differences in auditory cortical temporal acuity measures.

## Results

### Malleus removal surgery at P11 in WT mice mimicked hearing loss in *Df1*/+ mice

Like human 22q11.2 deletion carriers, *Df1*/+ mice exhibit a complex hearing phenotype with high inter-individual and even inter-ear variability (13,14). More than half of *Df1/+* mice have mild to moderate hearing loss in one or both ears (14,15,17). Hearing loss in *Df1*/+ mice arises primarily from increased risk for otitis media (middle-ear inflammation) as a result of hypoplastic Eustachian tube muscles which reduce the patency of the Eustachian tube (18). Inflammation and effusion in the middle ear disrupt sound-evoked vibration of the ossicles, causing conductive hearing loss (Figure 1B; compare to Figure 1A showing normal ear). In affected *Df1/+* mice, hearing loss is typically evident soon after ear opening at P11 and persists through adulthood (17).

**Figure 1.**
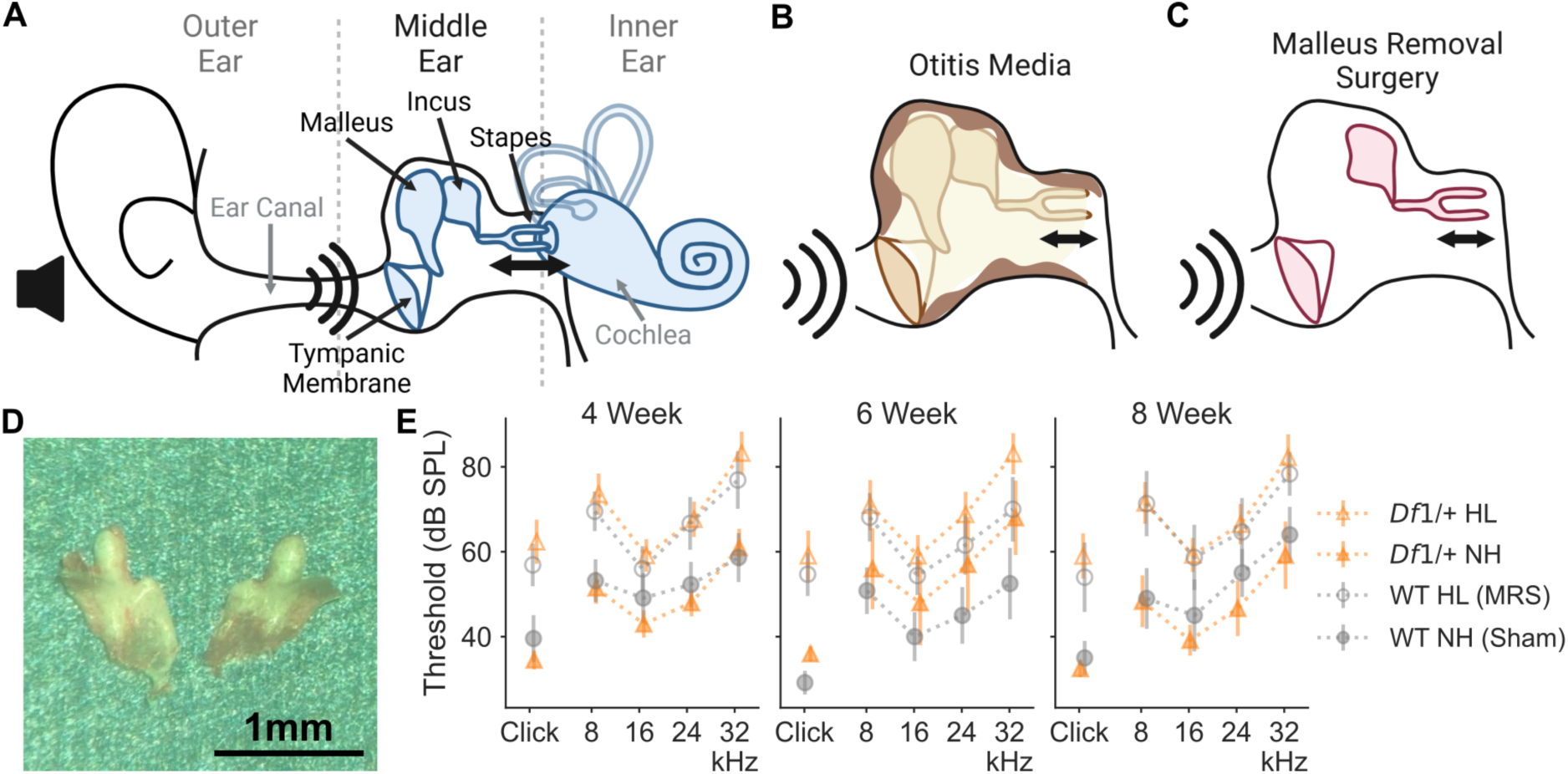
Malleus removal surgery in P11 WT mice mimics early conductive hearing loss observed in *Df1*/+ mice. **(A)** Anatomy of the mouse ear. The ossicular chain in the middle ear transmits sound vibrations to the inner ear via the connection of the malleus, incus, and stapes. **(B)** Hearing loss in *Df1/+* mice is primarily conductive, resulting from otitis media (inflammation and effusion in the middle ear) which disrupts ossicle movement and sound transmission. **(C)** Malleus removal surgery also disrupts sound transmission through the middle ear to produce conductive hearing loss. **(D)** Malleus bones dissected from a P11 WT mouse. **(E)** Click-evoked auditory brainstem response (ABR) thresholds show that malleus removal surgery in P11 WT mice caused conductive hearing loss similar to that observed in *Df1/+* mice. Orange dashed line, *Df1/+* mice with hearing loss (*Df1*/+ HL); orange solid line, *Df1/+* mice with normal hearing (*Df1*/+ NH); gray dashed line, WT mice with hearing loss (WT HL); gray solid line, WT mice with normal hearing (WT NH). Error bars represent mean ± 2 SEM.

To mimic a similarly early-onset and persistent conductive hearing loss in WT mice, we removed the malleus bone in P11 WT mouse pups to disrupt sound transduction (Figure 1C-D) (19,20). To evaluate the severity of hearing loss caused by malleus removal surgery (MRS), we measured click-evoked auditory brainstem responses (ABRs) in separate cohort of 32 mice, including *Df1*/+ mice (n = 15), WT mice with MRS (WT HL, n = 11) and WT mice with sham surgery (WT NH, n = 6) during development (Figure 1E). We classified an ear as having hearing loss (HL) if the click-evoked ABR was higher than 42 dB SPL (2.5 standard deviations above the mean estimated from hearing thresholds of WT ears with sham surgery); otherwise, the ear was classified as normal hearing (NH). By definition, then, ears in *Df1*/+ HL and WT HL groups had elevated hearing thresholds compared to those in *Df1*/+ NH and WT NH groups (Week 8: *Df1*/+ HL 59.00 ± 2.73 dB SPL, *Df1*/+ NH 32.50 ± 1.12 dB SPL, WT HL 54.00 ± 4.23 dB SPL, WT NH 35.00 ± 2.24 dB SPL; see Supplementary Table 1 for hearing thresholds at other ages). No significant differences in hearing thresholds were seen between *Df1*/+ HL and WT HL groups, nor between *Df1*/+ NH and WT NH groups (Week 8, Kruskal-Wallis test, H(_3, 46_) = 24.70, p < 0.001, post-hoc, p*_Df1_*_/+ HL - *Df1*/NH_ < 0.001, p_WT-HL - WT NH_ = 0.03, p*_Df1_*_/+ HL - WT HL_ = 0.43, p*_Df1_*_/+ NH - WT NH_ = 0.75). Tone-evoked ABRs revealed similarly increased thresholds across frequencies, consistent with the expected features of conductive hearing loss (Figure 1E). In summary, malleus removal surgery or sham surgery at P11 in WT mice produced hearing phenotypes in WT HL and WT NH mice comparable to those observed in *Df1*/+ HL and *Df1/+* NH mice, respectively.

### Intracranial recording of single-unit responses to gap-in-noise stimuli reveals effects of genotype and hearing phenotype on neural sensitivity to noise transients

We then studied the effects of the *Df1*/+ deletion and hearing loss on neuronal responses in the auditory cortex, focusing first on comparisons between WT mice with normal hearing, *Df1/+* mice with normal hearing, and WT mice with hearing loss. For analyses reported here, we used the subset of animals with equivalent hearing thresholds in the two ears or better hearing in the left ear, which was contralateral to the hemisphere used for cortical recording (Figure 2A; *Df1*/+ NH: *Df1*/+ mice with both ears normal hearing, n = 5; WT HL: WT mice with malleus removal surgery in both ears, n = 6; WT NH: WT mice with sham surgery in both ears, n = 10). Neural responses were recorded from the right primary auditory cortex using Neuropixels 1.0 probes while the awake mice passively listened to gap-in-noise stimuli (Figure 2B-C).

**Figure 2.**
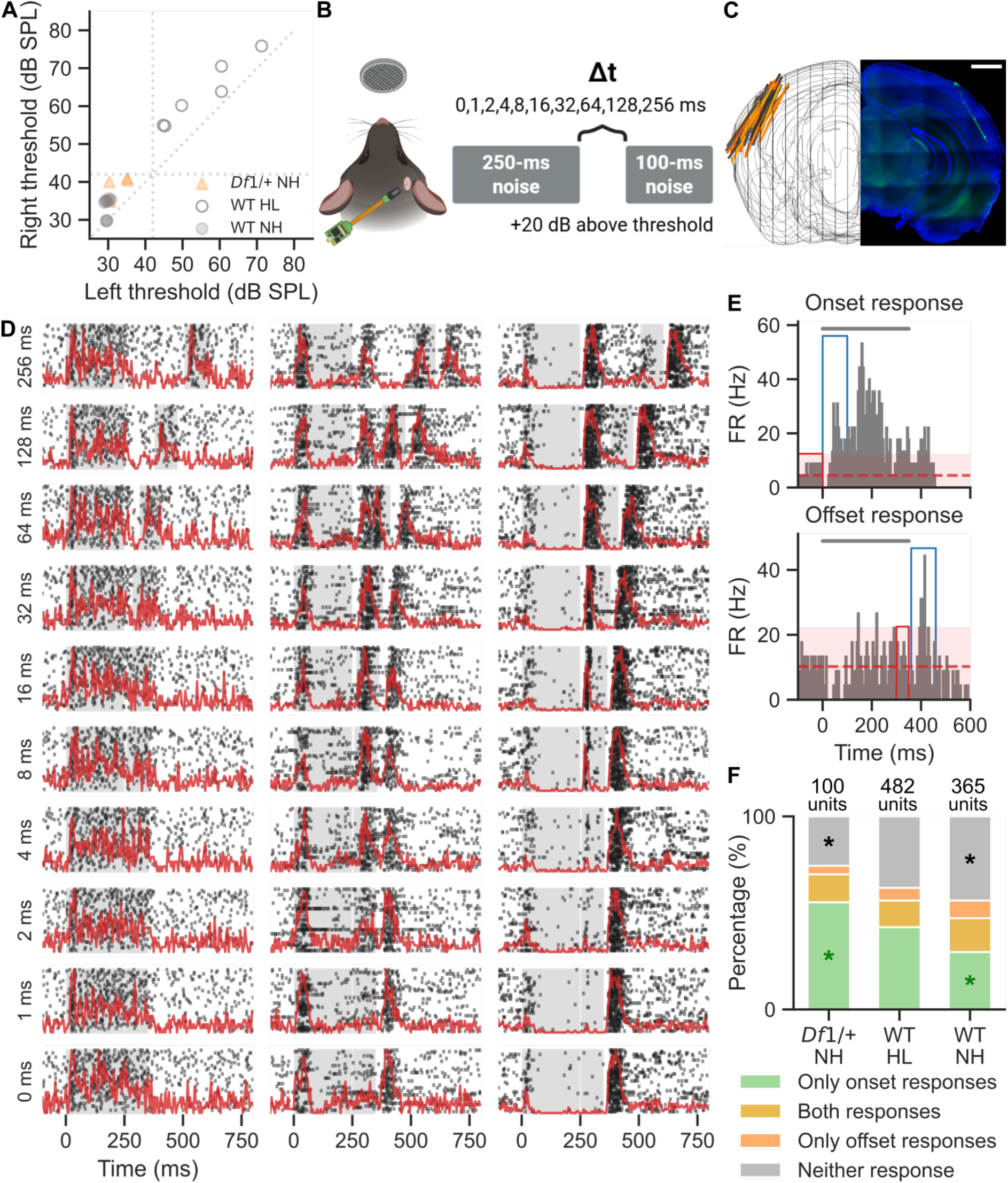
Neuropixels recordings reveal diverse neuronal responses to gap-in-noise stimuli in the auditory cortex of awake mice, and differences between animal groups in neuronal sensitivity to sound onsets and offsets. **(A)** Hearing thresholds were measured in each ear of each mouse prior to auditory cortical recording. We focused first on comparing *Df1*/+ NH (filled orange triangle, n = 5), WT HL (unfilled grey circle, n = 6) and WT NH (filled grey circle, n=10) mice. Mice with worse hearing in the left ear (not shown) were excluded from the study, to avoid understimulating the right (contralateral) auditory cortex where neuronal responses were recorded. **(B)** Single-unit activities were recorded acutely using Neuropixels 1.0 probes in the right auditory cortex of awake passively listening mice presented with gap-in-noise stimuli of varying gap durations. Noise intensity was individually adjusted for each mouse to be 20 dB above the left ear hearing threshold (which was the better-hearing or equivalent-hearing ear for all mice). **(C)** Probe traces in histological sections (right) and reconstructed trajectories through the auditory cortex (left). Right: green, DiO; blue, DAPI; scale bar, 1 mm. **(D)** Raster plots and peri-stimulus time histograms (PSTHs) show neuronal responses to gap-in-noise stimuli from three example units. Gray area, noise presentation; black dots, spike events; red line, PSTH. **(E)** Definition of onset and offset responses in single units. The gray histogram represents neural activity surrounding noise presentation (gray bar above). Transient responses were defined as at least two consecutive bins with significant increases in firing rate within the analysis window (blue) relative to the reference window (cyan; nonsignificant range shown in red). To capture suppression in neural firing by sound, significantly reduced responses were included for onset response detection (see Methods). **(F)** Distributions of units with different forms of sensitivity to noise onset and offset transients revealed significant differences between groups, particularly between *Df1*/+ NH and WT NH mice (marked with *) for units with onset-only responses (green) and units with no transient responses (gray).

Noise levels were set to 20 dB above the threshold of the better hearing ear to prevent overstimulation of the animals. Gap-in-noise stimuli consisted of a first 250-ms noise and a second 100-ms noise separated by a silent gap with duration of 0, 1, 2, 4, 8, 16, 32, 64, 128 or 256 ms. Primary auditory units showed various responses to the gap-in-noise stimuli, including transient responses to noise onset or offset and sustained responses to noise (Figure 2D). These transient responses to the offset of the first noise and/or the onset of the second noise gradually emerged with increasing gap durations, indicating the detection of the gap.

The proportion of units with transient responses to noise onset and/or offset varied between the three animal groups (Figure 2E). All units that passed the physiological criteria for primary auditory units (see Supplementary Methods) were clustered into units with only onset responses, units with only offset responses, units with both responses and units without transient responses (Figure 2F). These four clusters were distributed differently between animal groups (χ^2^_(6, 947)_ = 32.15, p < 0.001, post-hoc tests, p *_Df1_*_/+ NH - WT NH_ <0.001, p _WT NH - WT HL_ = 0.0014). *Df1*/+ NH and WT HL mice had a higher percentage of units with only onset responses and WT NH mice had a lower percentage of those units. Meanwhile, WT NH mice had the highest percentage of units that did not respond to either noise onset or offset. Thus, proportions of auditory responsive neurons with sensitivity to sound onsets and offsets depended on both genotype and hearing phenotype.

### Robust measurement of single-unit gap duration thresholds shows both hearing loss and 22q11.2 deletion are associated with temporal processing deficits

At the single-unit level, gap duration thresholds (GDTs) are often measured purely based on the onset responses to the second noise, ignoring the potential contribution of the offset responses to the first noise (21,22). Here, we propose a new method for analyzing GDTs at the single-unit level that utilizes all transient and other responses within and immediately following the gap. This method is general and robust enough that it can easily be applied to analysis of temporal acuity in other brain areas and sensory systems.

GDTs for single units were measured by comparing the deviance of neural responses around the gap in different gap conditions to the sustained responses in the 0-ms (no gap) condition (Figure 3A-B; see Methods and Supplementary Figure 1 for more details). The analysis period for gap detection was defined as extending from the start of the gap to the end of the second noise. We calculated the mean and standard deviation over time in bin-by-bin firing rates in this analysis period for the 0-ms gap condition (i.e., last 100 ms of the 350 ms continuous noise), and the root-mean-squared deviance (RMSD) over time from this mean for the analysis period in other gap conditions. Since the RMSD would be expected to rise rapidly and then level off as gap duration increased, a sigmoidal function was fitted to the RMSD values as a function of gap duration. The GDT for a unit could then be robustly defined as the gap duration at which the sigmoidal fit exceeded 2 times the standard deviation over time of the neural responses in analysis period for the 0-ms (no gap) condition, thereby taking into account that the variance in the neuron’s response to sustained noise affects the reliability of gap detection at the single-unit level.

**Figure 3.**
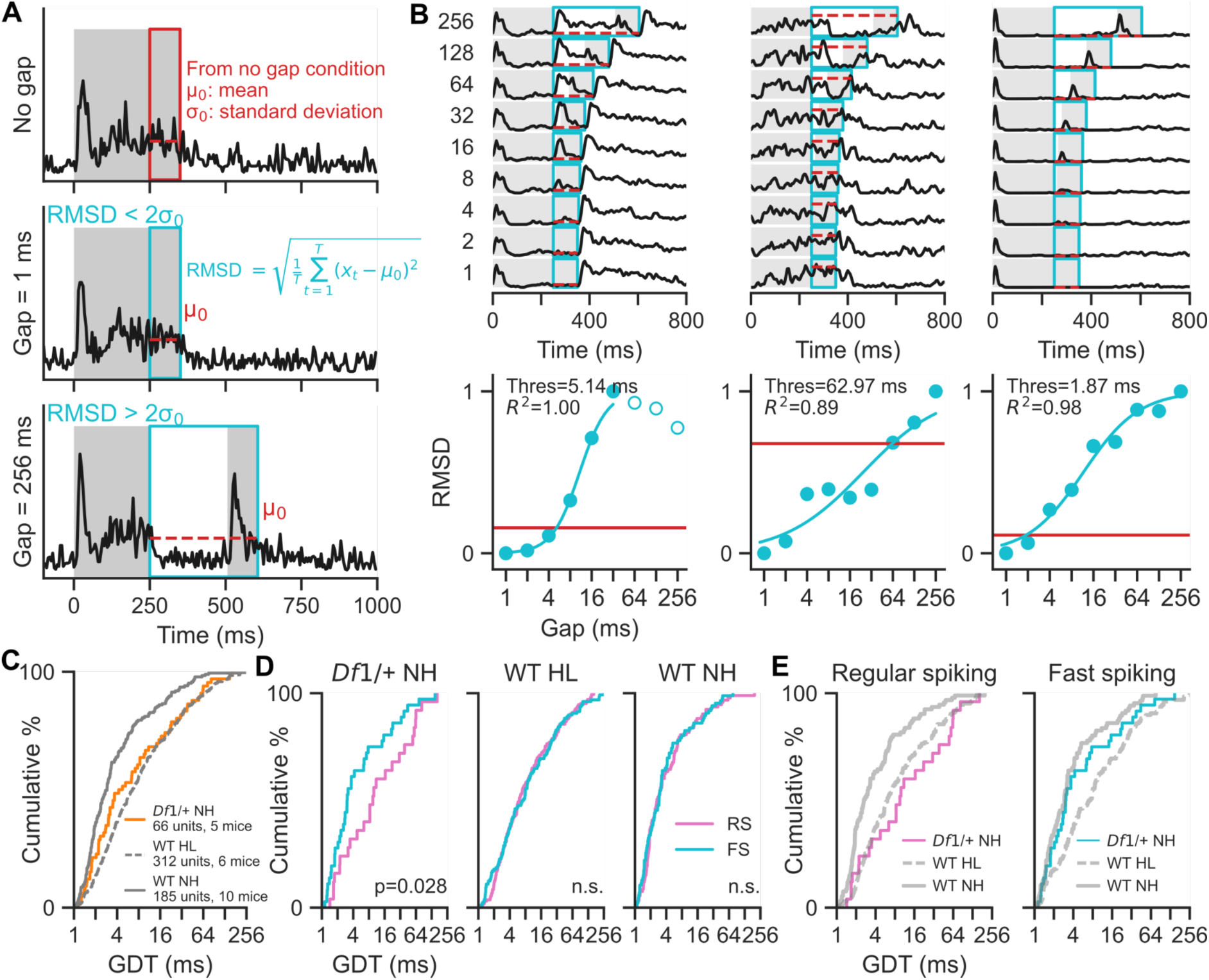
Hearing loss and the *Df1/+* deletion both elevated single-unit gap duration thresholds (GDTs), but effects of the *Df1/+* deletion were specific to regular-spiking units. (A) Single-unit GDT was estimated by comparing the root mean squared deviation (RMSD) of neuronal responses in any gap condition to the no-gap condition, over a time period extending from the end of the 250 ms first noise segment to the end of the 100 ms second noise segment. For the no-gap condition (top, red box) this period was a continuation of noise without a silent gap. For gap conditions (middle and bottom, cyan boxes), this period included a silent gap of variable duration. **(B)** Example GDT measurement in three example units. The RMSD of responses to the gap (highlighted with cyan shaded area in the top) was normalized (cyan dots in the bottom) and then fitted with a sigmoid function (cyan curve). The gap duration that corresponded to an RMSD 2 times as large as the standard deviation for the no-gap condition (red line) was defined as the GDT for that unit. Hollow circles represent data points not included in the sigmoid fitting. **(C)** Cumulative distribution functions of GDT for all units. GDTs were longer in units from WT HL mice than WT NH mice. GDTs were also longer in units from *Df1/+* NH mice than WT NH mice. **(D)** Cumulative distribution functions of GDT for units that could be classified with over 95% confidence as regular-spiking (RS, pink) or fast-spiking (FS, cyan) units. GDTs of RS units were significantly longer than GDTs of FS units in *Df1/+* NH mice. No significant differences were evident between GDTs of RS and FS units in either WT HL or WT NH mice. **(E)** For RS units (left), GDTs were longer in *Df1/+* NH mice than in WT NH mice. In contrast, for FS units (right), GDTs in *Df1/+* NH mice were similar to FS unit GDTs in WT NH mice.

This analysis revealed that GDTs of auditory cortical neurons in WT HL mice were the longest of any mouse group and significantly different from those in WT NH mice, indicating effects of hearing loss. Meanwhile, GDTs of units recorded in *Df1/+* NH mice were mildly elevated compared to those in WT NH mice, revealing effects of the 22q11.2 deletion (Kruskal-Wallis test, H_(2, 563)_ = 45.94, p < 0.001, post-hoc tests, p*_Df1_*_/+ NH - WT NH_ = 0.0057, p_WT HL-WT NH_ < 0.001; Figure 3C). Thus, single-unit measures of GDTs revealed that temporal acuity of neurons in the auditory cortex was dependent on both the hearing phenotype and the genotype of the mouse. In further analysis of units with onset and/or offset responses, we found that in normal hearing mice (both WT NH and *Df1/+* NH groups), units with only-onset or both onset and offset responses had shorter GDTs than units with only-offset responses (Supplementary Figure 2).

We also separated neurons based on their waveform features into putative excitatory neurons (regular-spiking units, RS) and putative parvalbumin-positive (PV^+^) interneurons (fast-spiking units, FS) (Supplementary Figure 3). Surprisingly, given evidence for higher temporal acuity in PV^+^ cells than in pyramidal neurons (22), we found no significant differences in GDT distributions between fast-spiking and regular-spiking in either of the WT groups (Mann-Whitney U test; WT HL, U_(166, 128)_ = 10679, p_WT HL_ = 0.94; WT NH, U_(93, 86)_ = 4185, p_WT NH_ = 0.59; Figure 3D), even though fast-spiking units were more sensitive to transient onset and offset events than regular-spiking units. In contrast, *Df1*/+ NH mice had divergent GDTs between RS and FS units (Mann-Whitney U test; *Df1*/+ NH, U_(25, 36)_ = 600, p *_Df1_*_/+ NH_ = 0.028). In *Df1*/+ NH mice, GDTs of RS units were abnormally elevated related to those in WT NH mice and similar to those in WT HL mice (Kruskal-Wallis test, H_(2, 146)_ = 26.65, p < 0.001, post-hoc tests, p*_Df1_*_/+ NH - WT HL_ = 1, p*_Df1_*_/+ NH - WT NH_ = 0.0012, p_WT HL-WT NH_ < 0.001), while GDTs of FS units were similar to those in WT NH mice (Kruskal-Wallis test, H_(2, 158)_ = 20,55, p < 0.001, post-hoc tests, p*_Df1_*_/+ NH - WT HL_ = 0.047, p*_Df1_*_/+ NH - WT NH_ = 0.34, p_WT HL-WT NH_ < 0.001).

### Novel measures of gap duration thresholds for neuronal population responses reveal only effects of hearing loss

Next we asked whether differences between animal groups in single-unit GDT distributions were also evident at the level of neuronal population dynamics. We applied principal component analysis (PCA) to reduce the dimensionality of the neuronal population and used the top three principal components (PCs) for representation (Figure 4A). The dynamics of PCs mainly revealed transient responses to noise onset and noise offset (Figure 4B). The most significant between-group differences appeared in PC1; responses to sustained noise were stronger in mice with hearing loss.

**Figure 4.**
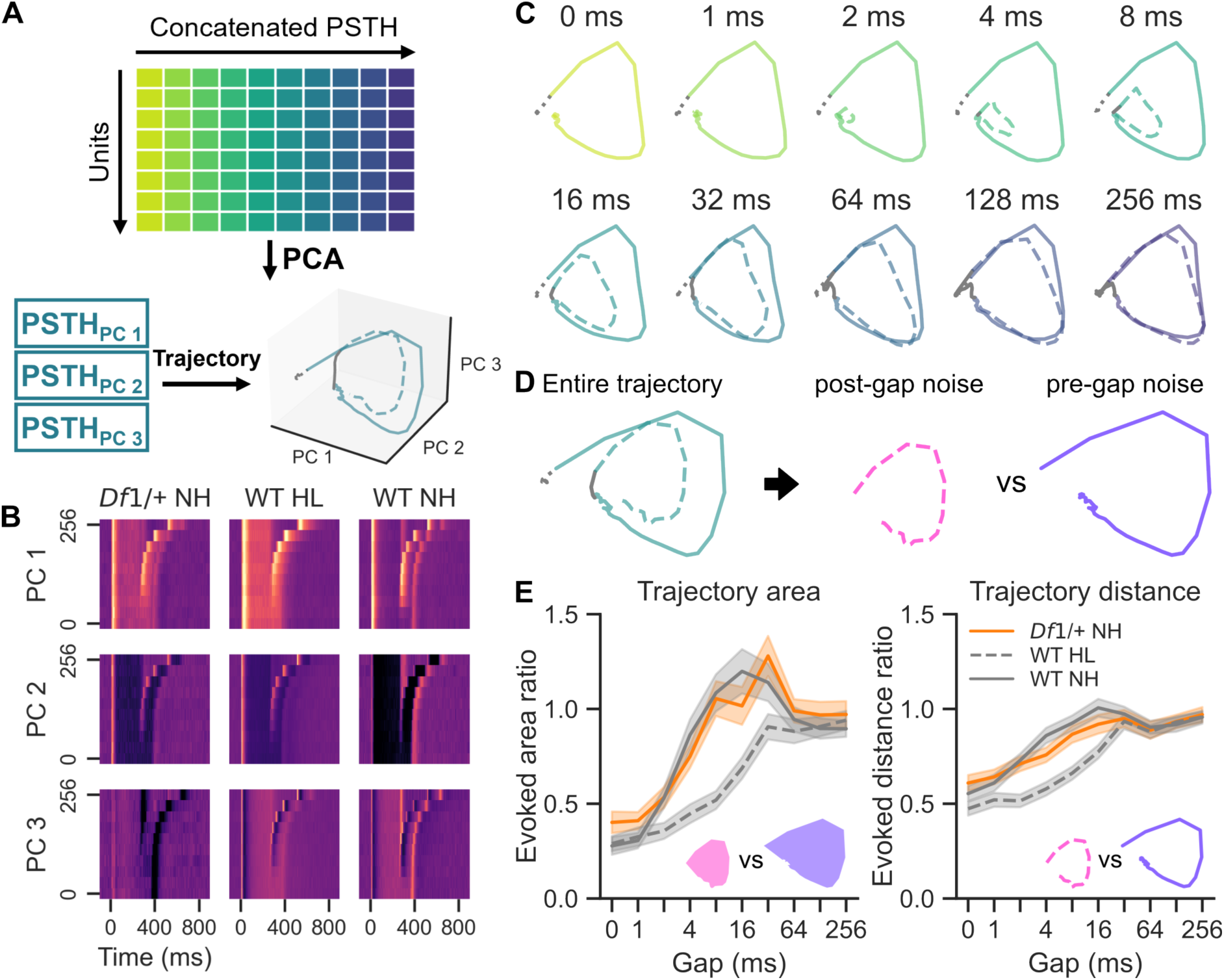
Analysis of neuronal population dynamics revealed elevated gap duration thresholds in mice with hearing loss. **(A)** Population activity was analyzed using principal component analysis. PSTHs from different gap duration conditions were concatenated for each neuron and then principal components (PCs) estimated to capture variability across neurons at each time point. Concatenated trial conditions were subsequently unwrapped for display of population trajectories in PC space for different gap conditions. **(B)** The temporal evolution of principal components captured population response dynamics across gap duration conditions in different animal groups. All PC weightings are shown on the same color scale. **(C)** In the space defined by the first three PCs in WT HL mice, trajectories for the second noise burst (colored dashed line) approached those for the first noise burst (colored solid line) as gap duration increased. The trajectory during the gap (solid gray line) was distinct from those during noise periods. **(D)** The area and distance ratios of trajectories evoked by the post-gap noise relative to those evoked by the pre-gap noise were calculated to compare gap-duration sensitivity of the neuronal population across groups. **(E)** Bootstrapped estimates of trajectory area (defined as the sum of triangular areas formed between adjacent timepoints and the baseline starting point) showed that population responses in WT HL mice were less sensitive to short gaps in noises than those in normal hearing mice. Bootstrapped estimates of trajectory distance (defined as the sum of distances between adjacent timepoints) revealed a similar trend toward reduced population sensitivity to short gaps in mice with hearing loss. Shaded areas represent mean ± 3.3 SEM, corresponding to the 99.9% confidence interval.

To better visualize the dynamics of population responses to gap-in-noise stimuli, we reconstructed the trajectory of neuronal responses to gap-in-noise stimuli in PC1-3 space (Figure 4C). When the noise was present, the trajectory moved outward quickly and then stabilized at an evoked location. Once the noise was off, the trajectory gradually returned to the starting point. When the second noise burst occurred, the trajectory rebounded. The extent of these rebound activities was dependent on the gap length; smaller gaps resulted in minimal rebound activity, whereas extended gaps led to increasingly pronounced rebound activities to the second noises.

We used the distance of the trajectory and the area covered by the trajectory to approximate the gap duration threshold of the neuronal populations recorded in each animal group (Figure 4D and 4E). The trajectory distance was calculated as the cumulative sum of the Euclidean distances between consecutive time points. The area covered by the trajectory was estimated by the cumulative sum of curved triangles formed along the trajectory. Both the trajectory distance and trajectory area indicated that at the neuronal population level, gap duration thresholds in mice with normal hearing were shorter compared to those with hearing loss, regardless of their genetic background. Thus, effects of hearing loss on auditory cortical temporal acuity dominated effects of genotype at the neuronal population level.

### Distinct effects of 22q11.2 deletion and hearing loss are evident even under comorbid conditions

Finally, to investigate the combined effects of 22q11.2 deletion and hearing loss on auditory temporal processing, we studied *Df1*/+ mice with naturally occurring hearing loss (*Df1*/+ HL). Like humans with the 22q11.2 deletion, *Df1/+* mice can have either normal or impaired hearing, and individuals with impaired hearing may have different levels of hearing loss in the two ea(14,18). To ensure adequate stimulation of the recorded right hemisphere, we included in our analysis only mice with better hearing in the left ear (contralateral to the recorded hemisphere), since stimuli were always adjusted in loudness relative to the better-ear hearing threshold. Applying this criterion reduced the eligible *Df1*/+ HL cohort from 7 to 2 animals, and the following results should therefore be interpreted as preliminary.

Despite the limited sample size, cortical recordings from *Df1*/+ HL mice displayed deficits consistent with both 22q11.2 deletion and hearing loss (Figure 5). As observed in *Df1*/+ NH mice but not WT mice, RS and FS units in *Df1*/+ HL mice showed divergent GDT profiles. Only RS units in *Df1/+* HL mice exhibited elevated GDTs comparable to those observed in both RS and FS units in WT HL mice. Additionally, and unlike *Df1/+* NH mice, *Df1*/+ HL mice exhibited abnormally slow growth in neural population activity trajectories for short gaps in noise, consistent with hearing loss-associated auditory temporal processing deficits observed in WT HL mice.

**Figure 5.**
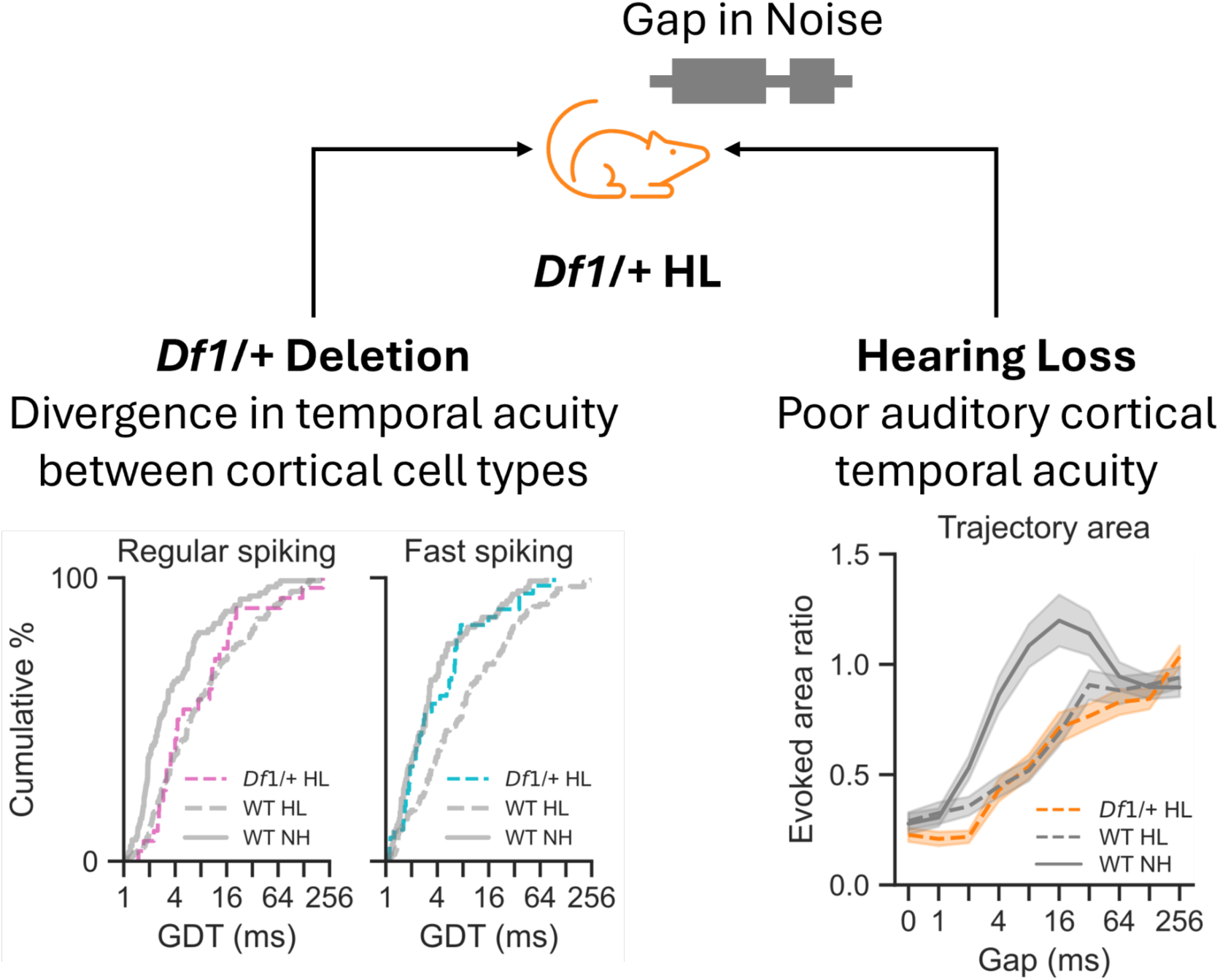
Distinct effects of *Df1*/+ deletion and hearing loss remain evident under comorbid conditions in *Df1*/+ HL mice. Like *Df1/+* NH mice (compare left panels here to Figure 3E), *Df1/+* HL mice showed divergence in temporal acuity between regular-spiking (putative excitatory) and fast-spiking (putative inhibitory) cortical cell types, relative to WT NH and WT HL mice. Like WT HL mice (compare right panel to Figure 4E), *Df1/+* HL mice showed poorer temporal acuity of neural population activity, with smaller excursions in auditory cortical population trajectories for brief silent gaps despite the fact that sound levels were adjusted relative to hearing threshold to ensure equivalent loudness. Thus, distinct effects of the *Df1/+* deletion and hearing loss were observed even under comorbid conditions. These results suggest that the 22q11.2 deletion and hearing loss disrupt auditory cortical temporal acuity through different mechanisms.

## Discussion

Auditory cortical dysfunction is commonly recognized as an endophenotype of psychosis (1), and provides a window for investigating how multiple risk factors contribute to the generation of neuropathology in psychiatric disease. In this study, we examined auditory cortical abnormalities associated with hearing loss and the 22q11.2 deletion using gap-in-noise stimuli as a sensitive index of temporal dynamics in the auditory cortex. We first showed that malleus removal surgery in WT mice induced conductive hearing loss comparable to that observed in a subset of *Df1/+* mice. We then established robust methods to quantify duration thresholds for responses to a brief gap in noise at both the single-unit and neuronal population levels, enabling precise assessment of auditory cortical temporal acuity across mice with different genetic backgrounds and hearing sensitivities. Finally, we deployed these methods to disentangle the effects of hearing loss and the *Df1/+* deletion on auditory cortical temporal acuity.

The results indicated that hearing loss and the *Df1/+* deletion alter temporal dynamics in the auditory cortex both independently and jointly. Hearing loss impaired auditory cortical temporal acuity at both the single-unit and neuronal population levels in WT mice, even though sound intensity was adjusted to be constant relative to hearing threshold for each mouse. In contrast, the *Df1/+* deletion alone caused a more modest disruption of cortical temporal acuity, and its effects were detectable only at the single-unit level. Importantly, the effects of the sensory and genetic risk factors differed in their cell-type specificity: hearing loss impaired gap duration thresholds in both putative excitatory and putative inhibitory neurons, whereas the *Df1/+* deletion selectively affected putative excitatory neurons. In a small number of *Df1/+* mice with naturally occurring hearing loss, we observed that distinct effects of the *Df1/+* deletion and hearing loss were preserved under comorbid conditions. Together, these observations suggest that hearing loss and the 22q11.2 deletion disrupt cortical temporal acuity through distinct neural mechanisms.

Our data align with previous behavioural and neurophysiological studies showing that hearing loss impairs gap detection (10). Young adults with normal hearing can detect silent gaps as short as 2-ms in continuous noise, and hearing loss elevates the detection thresholds by a factor of two or more. Similar gap detection threshold shifts caused by hearing loss have also been reported in animal models (21,23). The differences we observed in gap duration thresholds between mouse groups with and without hearing loss were similar in magnitude to those reported in previous behavioural and neurophysiological studies in rodents. In contrast to a report that PV^+^ inhibitory neurons have shorter gap duration threshold (22), in WT mice no clear differences in gap duration thresholds were observed between fast-spiking units and regular spiking units, and hearing loss had a similar impact on these two neuron subtypes.

The influence of the 22q11.2 deletion on auditory cortical temporal processing can be summarised in three key observations. First, at the single-unit level, gap duration thresholds were mildly elevated even in *Df1*/+ mice with normal hearing, relative to WT mice with normal hearing. This mild impairment of cortical temporal acuity in *Df1/+* mice might arise from synaptic changes caused by haploinsufficiency of genes located in the deletion region (24,25). Second, an unusually high proportion of auditory cortical units in *Df1*/+ mice with normal hearing had onset-only responses to noise bursts. Since units with onset-only responses also tended to have the shortest gap duration thresholds, this increase in the proportion of cells with onset-only transient responses seems more likely to be a compensating than contributing factor for auditory temporal processing deficits. Third, and most strikingly, the *Df1/+* deletion was associated with divergence in temporal acuity between neuron subtypes: gap duration thresholds were longer in regular-spiking than fast-spiking units in *Df1/+* mice with or without hearing loss. Longer gap duration thresholds in regular-spiking units (putative pyramidal excitatory neurons) may be related to reduced evoked excitability (26) and disrupted thalamocortical connection (27) in mouse models of 22q11.2 Deletion Syndrome. Fast-spiking units (putative PV^+^ inhibitory neurons) appeared more robust in *Df1/+* mice, with shorter gap duration thresholds that were similar to those observed in WT mice with normal hearing and minimally affected by hearing loss.

In conclusion, this work illustrates how animal models can be used to address questions about the role of comorbidities in psychiatric disease that are difficult or impractical to address in humans. In humans with the 22q11.2 deletion, it is not known whether comorbid hearing loss exacerbates risk of developing psychiatric disease, although hearing loss is known to be a risk factor for psychosis in the general population. In mice, we were able to compare genetically homogeneous groups of mice with and without hearing loss and with and without a mouse homologue of the 22q11.2 deletion, to disentangle effects of sensory and genetic risk factors with a precision that would be impossible to achieve even in studies of human siblings or twins. We analyzed effects of these risk factors on a sensitive measure of auditory brain function: cortical temporal dynamics evoked by brief silent gaps in noise. Results showed that the 22q11.2 deletion alone alters auditory cortical temporal processing with cell-type selectivity, but effects of hearing loss are more profound at both the single-unit and neuronal population levels. Thus, these findings provide a mechanistic proof-of-concept that auditory cortical dysfunction in individuals with genetic risk for psychiatric disease may manifest differently with and without hearing loss comorbidity.

## Acknowledgements

We thank Eleanor Benoit for her assistance with data collection. We are grateful to Dan Sanes for providing detailed advice on malleus removal surgery in mice. We also thank Elvira Bramon, Peter Keating, and Nick Lesica for their contributions to supervision of Chen Lu’s PhD research. This work was supported by the UCL Institute for Mental Health (Small Grant 2022 to JFL) and the UK Medical Research Council (grant MR/P006221/1 to JFL), and also benefitted from technical support funded by the NIHR UCLH Biomedical Research Centre Hearing Health Theme.

## Author contributions

**CL**: conceptualization, methodology, software, investigation, formal analysis, visualization, writing.

**JFL**: conceptualization, resources, writing, supervision, funding acquisition.

## Data Availability

Raw data and/or analysis code are available from the authors upon request.

## Supplementary Methods

### Animals

*Df1/*+ mice were originally developed from a 129SvEvBrd X C57BL/6J background (1) and had been maintained on a C57BL/6J background for well over 25 generations, through pairing of *Df1/+* males either with WT females from the colony or (at least yearly) with newly acquired C57BL/6J females from Charles River UK. WT mice used in this study were also C57BL/6J mice purchased from Charles River UK. Mice of both sexes were included in all parts of this study. Mice were raised in standard cages and mouse housing facilities, on a standard 12 h-light/12 h-dark cycle. All experiments were performed in accordance with a Home Office project licence approved under the United Kingdom Animal Scientific Procedures Act of 1986.

### Malleus removal surgery

Approximately half of *Df1/+* mice experience chronic hearing impairment in one or both ears, arising from early-onset middle ear inflammation which causes conductive hearing loss (2,3). To generate an early-onset and comparable conductive hearing loss in a subset of WT animals, we performed malleus removal in WT mouse pups at P11, before the opening of the ear canal (4,5). Pups were anesthetized with isoflurane (3% for induction, 1.5-2% for maintenance with 2 L/min O_2_) and body temperature was maintained at 37 °C using a homeostatic heat pad. Carprofen (5 mg/kg body weight) was administered subcutaneously for analgesia. A postauricular skin incision was made and the cartilage forming the ear canal was cut through to expose the tympanic membrane. After piercing the tympanic membrane, the malleus was removed by grasping the orbicular apophysis through this opening using a pair of fine forceps. The skin was then closed with Vetbond Tissue Adhesive (3M). Once recovered from anesthesia in a warm cage (30-35 °C), the pups were returned to the nest together with their siblings. For sham surgeries, every step was performed identically except that the tympanic membrane was left intact and the malleus was not removed.

### Auditory brainstem response recording

The setup for auditory brainstem response recording was the same as described previously (6).

Auditory stimuli were generated at a sample rate of 195,312.5 Hz using a digital signal processor (Tucker Davis Technologies, TDT RX6). The signals were attenuated (TDT PA5), amplified (TDT SA1), and delivered via a free-field speaker (TDT FF1) positioned 10 cm from the tested ear. The speaker output was calibrated to within ± 1.5 dB of the target values (100 or 105 dB SPL) using a free-field 1/4” microphone (G.R.A.S.) placed at the location of the stimulated ear.

Auditory brainstem response (ABR) signals were recorded using subdermal electrodes (Rochester Medical) positioned at the vertex (positive electrode), the bulla of the tested ear (reference electrode), and the olfactory bulb (ground electrode). Data were acquired at a sample rate of 24,414 Hz (TDT RX5) using a low-impedance headstage and signal amplifier (TDT RA4LI and RA16SD, with an overall gain of 20x, and 2.2 Hz - 7.5 kHz filtering), supplemented by a custom low-pass filter designed to remove attenuation switching transients (cutoff at 100 kHz). Stimulus presentation and data acquisition were managed using TDT BrainWare software in combination with custom MATLAB scripts.

Animals were anesthetized with a cocktail solution of ketamine (75 mg/kg body weight) and medetomidine (0.3 mg/kg body weight), administered via intraperitoneal injection. Carprofen (5 mg/kg body weight) was administered subcutaneously for analgesia. Saline or Hartmann’s solution (0.1 mL) was administered for hydration. To protect the eyes during the procedure, a liquid gel (Viscotears or GelTears) was applied. The skin was disinfected with 70% ethanol. The body temperature of each mouse was maintained at 37 °C using a homeostatic blanket. Breathing rate and toe-pinch response were monitored frequently to ensure stable anaesthesia. For recovery ABR measurements, animals were administered atipamezole (2 mg/kg body weight) and placed in a warm cage (26-30 °C) until they fully recovered.

The ABR signal was measured as the potential difference between the filtered signals from the vertex electrode and the ear bulla electrode. The signals were filtered using a fifth-order Butterworth band-pass filter with a frequency range of 500–3000 Hz. Averaged ABR signals across different intensity levels were plotted in a stacked format. The hearing threshold was defined as the lowest sound intensity level that evoked an ABR response with at least one deflection larger than 2 standard errors in the pre-stimulus ABR signal.

### Intracranial recording with Neuropixels 1.0 probes

#### Acoustic stimuli

Auditory stimuli were generated using customised software written in MATLAB. Stimuli were converted to an analog signal using an external sound interface (RME Fireface UC), amplified via an amplifier (ROTEL), and played through a speaker (Peerless by Tymphany). The speaker was positioned 10 cm in front of the animal. The speaker output was calibrated to within ± 0.2 dB of the maximal calibration values (100 dB SPL for 8–64 kHz tones, 105 dB SPL for 16 kHz tone) using a free-field 1/4” microphone (G.R.A.S.), positioned at the location of the mouse’s head.

We used neural responses to pure tones varying in frequency and sound intensity to identify likely primary auditory cortical units. Pure tone frequencies ranged from 8 to 64 kHz, with 8 steps per octave. Pure tone sound intensity was adjusted based on the better-ear hearing threshold of the mouse and ranged from -20 to 30 dB HL in 5 dB increments relative to the hearing threshold. When the hearing thresholds differed between ears, the threshold of the better-hearing ear was used to avoid overstimulation. Each tone was played for 50 ms, with 5-ms onset and offset ramps, and the inter-stimulus interval was 500 ms. Different tones were presented in a pseudo-random order with 3 repetitions.

The noise level for the gap-in-noise stimulus was also adjusted for each mouse, to be 20 dB above the better-ear hearing threshold. The gap-in-noise stimulus was composed of a pair of white noise bursts, with a gap duration in the middle varying from 0 to 256 ms (0, 1, 2, 4, 8, 16, 32, 64, 128, 256 ms). The white noise was synthesized by summing discrete pre-calibrated pure tones from 8 to 64 kHz, spaced at 8 steps per octave. The first noise burst lasted for 250 ms, followed by a gap, and the second noise burst lasted for 100 ms. To avoid blurring the broadband transients, no ramping was used. The inter-trial interval was 1000 ms, and each gap had 45 repetitions. The trial order was pseudo-randomized.

#### Animal procedures

Adult mice (less than 12 weeks of age) were anesthetized with isoflurane (3% for inducting and 1.25-1.75% for maintaining with 2 L/min O_2_). An incision was made along the midline to expose the skull. The craniotomy site was marked for acute recording (AP -2.2 to -3.6 mm, ML 2.5 to 3.2 mm from Bregma). A small hole was drilled above the left frontal lobe, and a bone screw connected to a male pin was inserted for grounding. The bone screw and the headpost were fixed onto the skull using dental adhesive resin cement (Super-bond, Sun Medical). A well was constructed around the pre-marked craniotomy site. Dental cement was used to reinforce the walls of the well and to protect the wire and the base of the pin. The well was sealed with silicone sealant (Kwik-Cast, WPI) to protect the exposed skull. Warm saline (0.1 mL) was administered subcutaneously for hydration before removing the mouse from anesthesia. Carprofen (0.02–0.025 mg/mL in drinking water) was provided for the first three days following surgery.

Mice were habituated gradually to increasing durations of head-fixation for at least 5 days before the acute recording day to reduce stress.

On the morning of the acute recording, a craniotomy was performed following the marked site to expose the brain tissue. Anaesthesia was induced and briefly maintained with isoflurane (3% for induction and 1.25–2% for maintenance with 2 L/min O_2_). Carprofen (5 mg/kg body weight) and dexamethasone (0.2 mg/kg body weight) were administered subcutaneously. A craniotomy well was drilled along the marked trace to expose the brain tissue. Saline and silicone sealant (Kwik-Cast, WPI) were used to protect the exposed brain tissue. Warm saline was administered for hydration before the mouse was removed from anaesthesia.

Once the mouse had recovered from anesthesia, it was head-fixed in the recording apparatus. The silicone sealant cap was removed, and the craniotomy site was filled with sterile saline. A Neuropixels 1.0 probe (IMEC) stained with fluorescent dye DiO (Invitrogen) was gradually inserted into the brain tissue. The probe was referenced to the implanted bone screw and grounded to the anti-vibration table. Neural signals were transmitted to the headstage (IMEC), then to the PXIe acquisition module (IMEC) installed in a PXI chassis (PXIe-1071, National Instruments) and finally to the host computer. The probe was gradually lowered to a depth of 2000–2500 μm at an extremely slow speed of 2–2.5 μm/s with a micromanipulator (Sensapex). The recording software SpikeGLX was opened to monitor the changes in the signal when lowering the probe. After 10-minute settling for the probe, the recording was then started. Once the recording was complete, the probe was retracted at a speed of 5 μm/s for the first 100 μm, and then at a speed of 50μm/s for the remaining distance. The craniotomy site was washed with sterile saline and covered with silicone sealant before returning the mouse to the cage in the warm incubator.

Typically, each mouse underwent two recording sessions on the day when the craniotomy was opened. The interval between two recording sessions was usually around 3 hours, during which time the mouse was returned to a standard cage in a quiet, warm environment. The probe was cleaned using 1% Tergazyme enzyme detergent in lukewarm double-distilled water (∼30 °C) and then rinsed and soaked in double-distilled water for re-use.

After recordings were completed, mice were euthanized using pentobarbital (0.2 – 0.3 mL of 20 mg/mL). Following transcardiac perfusion with phosphate-buffered saline and subsequently with 4% paraformaldehyde, the brain tissue was harvested and soaked in 4% paraformaldehyde for an additional 24 hours. The brain tissue was sectioned into 50 μm slices using a vibratome (Leica VT1200S). Sections were mounted with DAPI Fluoromount-G (Invitrogen) and imaged using a widefield microscope (Zeiss Axio Imager 2).

### Data processing

Preprocessing was performed using the SpikeInterface (7). Action potential recordings were initially corrected for phase shifts and band-pass filtered (300–9000 Hz). Bad channels, which included damaged recording sites or sites located outside of the brain, were identified and removed. The signals were then re-referenced using a global common median reference to eliminate common noise. Spiking events were extracted from the signals using Kilosort 2.5 (8). A quality matrix was calculated to exclude invalid units based on the waveform features of units: amplitude median was set above 50 μV, amplitude cutoff was set below 0.25, inter-spike interval violation was set below 0.2, and isolation distance was above 15.

Putative fast-spiking units and regular-spiking units were clustered based on the peak-trough intervals and peak-trough ratios (9,10). Units classified as positive (with peak amplitude higher than trough amplitude) were excluded from further analysis (11). A Gaussian mixture model was applied and only units with clustering probabilities greater than 0.95 were retained.

Likely primary auditory units were defined based on physiological features: significant, short-latency tone-evoked responses. Pure tone trials involving all tones presented at intensity levels above the animal’s hearing threshold were pooled, and firing rates were compared between the 50-ms periods before and after tone onset to identify units with significant tone-evoked responses (Wilcoxon test, p<0.01). Then, the same pooled trials were then used to construct a post-stimulus time histogram (PSTH) with 2-ms bins, to identify the earliest bin following sound onset in which firing rate exceeded 3 standard deviations around the mean spontaneous firing rate in the 50-ms period before sound onset. If this latency was less than 50 ms, then the tone-responsive unit was defined as a likely primary (core) auditory cortical unit.

### Data analysis

Animals were grouped based on their genotypes and hearing sensitivities in both ears, as measured with click-evoked ABR at week 8 (*Df1*/+ HL: *Df1*/+ mice with both ears having hearing loss; *Df1*/+ NH: *Df1*/+ mice with both ears normal hearing; WT HL: WT mice with malleus removal surgery in both ears; WT NH: WT mice with sham surgery in both ears). Since sound levels for all stimuli were adjusted to be constant relative to the better-ear hearing threshold for each mouse, and since neural recordings were made in the right auditory cortex (which is most strongly driven by the left ear), we focused our analysis on animals for which the left ear was the better-hearing ear.

Statistical tests are defined in the text when used and were conducted two-tailed with ɑ=0.05 unless noted otherwise. Post-hoc pairwise tests were conducted with Holm-Bonferroni correction. Analysis of onset and offset responses, single-unit gap-duration thresholds, and population-level gap-duration thresholds are described briefly below and more extensively in the Results section.

---

**Supplementary Figure 1.**
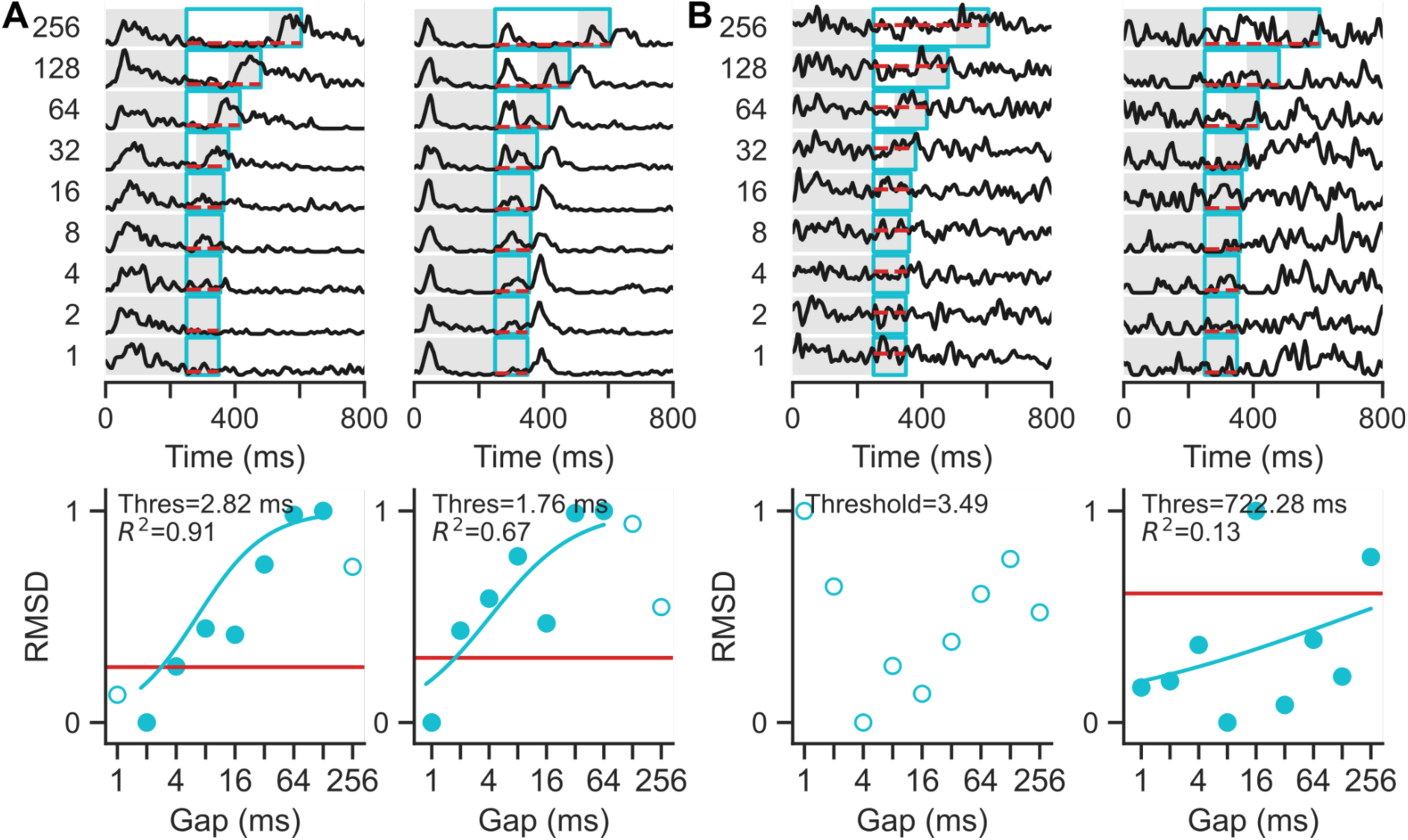
Additional examples of single-unit gap duration threshold (GDT) analysis methods. **(A)** Example units for which data from only a subset of gap durations were included in the sigmoid fit. When RMSD values decreased at longer gap durations (left) or were close to zero for shorter gaps (left and right), the sigmoid fit was performed after excluding these problematic data points (hollow circles). If the sigmoid fit was improved by the data exclusion, then the gap duration thresholds were estimated using this modified sigmoid fit. **(B)** Example units for which no gap detection thresholds could be identified.

**Supplementary Figure 2.**
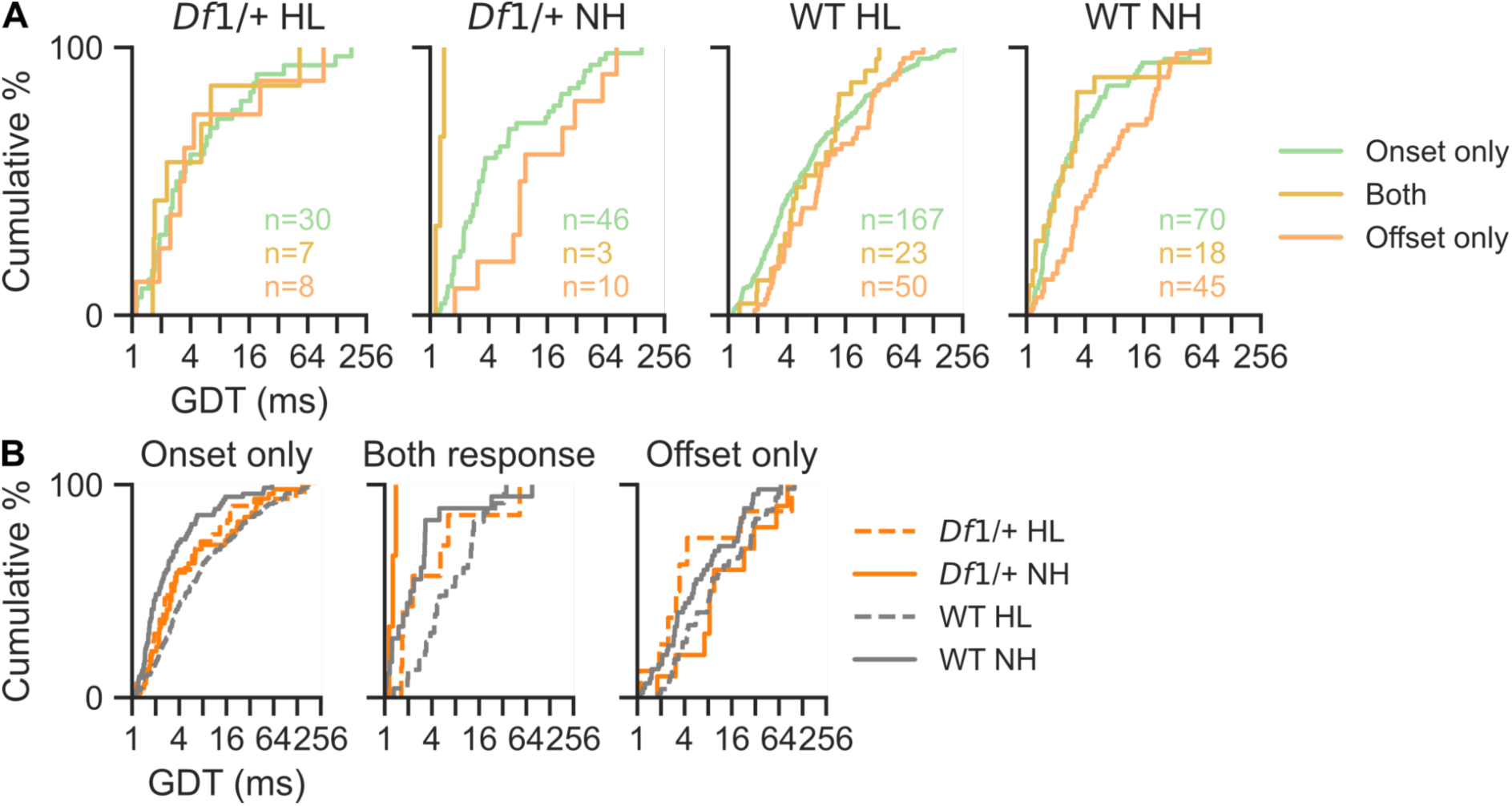
Distribution of gap duration thresholds (GDTs) in units with different onset and offset responses. **(A)** In normal hearing mice (*Df1/+* NH and WT NH), units with only offset responses had longer GDTs compared to units with only onset responses or with both responses (two-sample Kolmogorov-Smirnov test, units with only offset versus units with only onset and units with both onset and offset, p*_Df1_*_/+ HL_= 0.97, p*_Df1_*_/+ NH_= 0.015, p_WT HL_= 0.073, p_WT NH_ = 0.001). **(B)** For comparison across animal groups, units with onset-only responses in WT NH mice had the shortest GDTs while those in WT HL mice had the longest (Kruskal-Wallis test, p < 0.001; significant p-values in post-hoc tests, p_WT HL-WT NH_ < 0.0083, p*_Df1_*_/+ NH-WT NH_ <0.001). Units with both onset and offset responses showed similar trends between WT NH and WT HL mice (Kruskal-Wallis test, p = 0.00124; significant p-value in post-hoc tests, p_WT HL-WT NH_ = 0.0015). Results for *Df1*/+ mice remained unclear due to the low unit counts. For units with only offset responses, the significance for between-group differences was borderline (Kruskal-Wallis test, p = 0.0502).

**Supplementary Figure 3.**
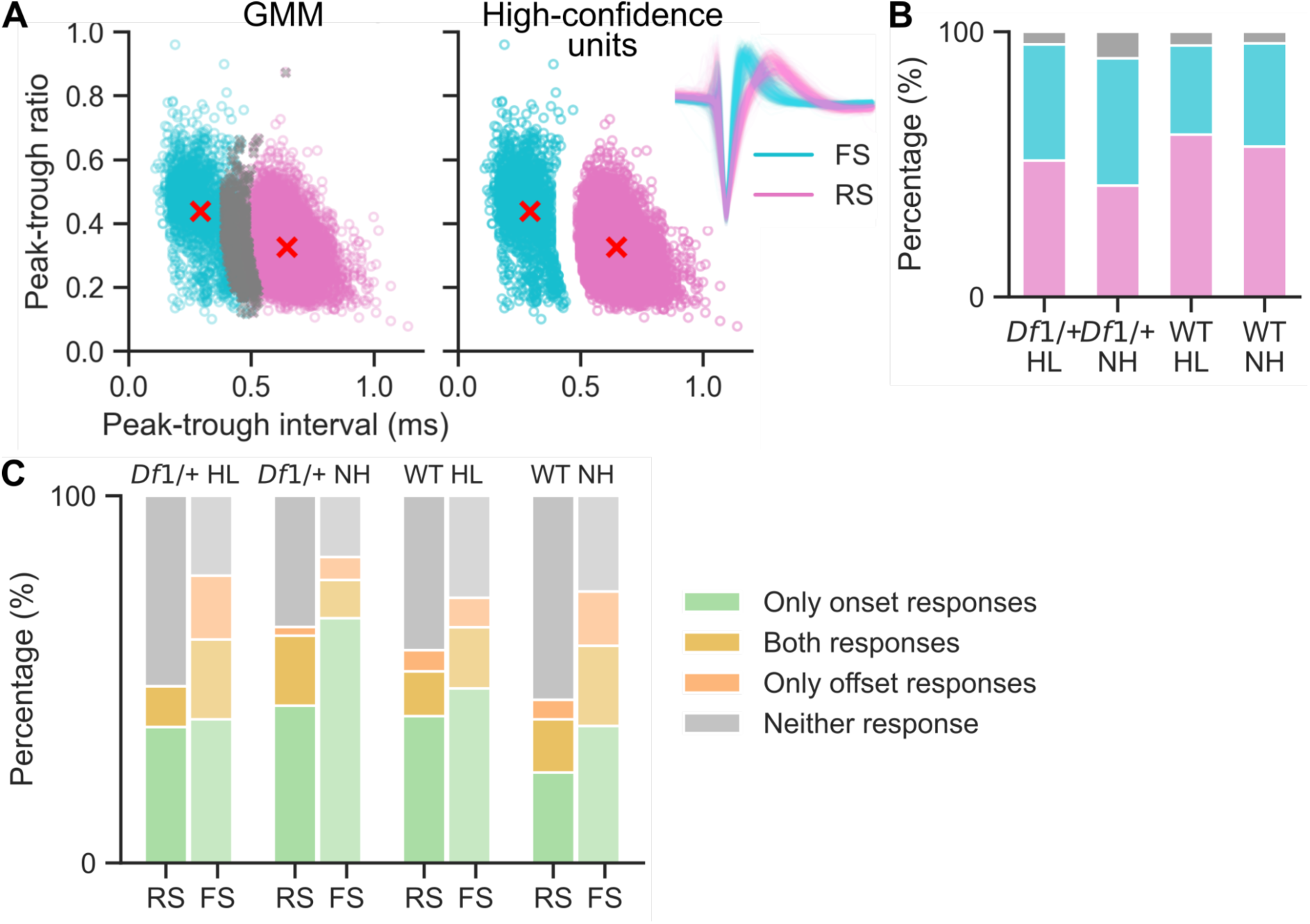
Classification of regular-spiking (RS) and fast-spiking (FS) units and their onset and offset response distributions. **(A)** After quantifying peak-trough interval and peak-trough ratio of spike waveform in every single unit, a Gaussian mixture model (GMM) was applied to cluster single units into regular-spiking and fast-spiking units. Units within the boundary area were defined as low confidence units and excluded. **(B)** The distribution of RS and FS units in recorded auditory units. **(C)** Fast-spiking units (putative PV+ interneurons) showed higher sensitivities to noise onset and offset transients compared to regular-spiking units (putative excitatory pyramidal neurons).

**Supplementary Table 1.** Hearing profiles in *Df1*/+ mice and WT mice during development. Values indicate mean ± standard error of ABR thresholds (dB SPL) measured in Weeks 4, 6 and 8 for animals in each group.

|  | Week 4 | Week 6 | Week 8 |
| --- | --- | --- | --- |
| *Df1/+* HL | Click: 62.35 ± 2.68<br>8kHz: 73.53 ± 2.53<br>16kHz: 59.12 ± 1.98<br>24kHz: 67.65 ± 2.15<br>32kHz: 83.24 ± 2.61 | Click: 59.05 ± 3.04<br>8kHz: 70.95 ± 3.04<br>16kHz: 59.05 ± 2.50<br>24kHz: 68.81 ± 2.71<br>32kHz: 83.10 ± 2.48 | Click: 59.00 ± 2.73<br>8kHz: 71.50 ± 2.52<br>16kHz: 59.25 ± 2.06<br>24kHz: 66.50 ± 2.44<br>32kHz: 82.25 ± 2.77 |
| *Df1/+* NH | Click: 34.50 ± 1.17<br>8kHz: 51.50 ± 1.98<br>16kHz: 43.00 ± 1.70<br>24kHz: 48.00 ± 1.70<br>32kHz: 61.00 ± 2.33 | Click: 36.00 ± 1.00<br>8kHz: 56.00 ± 5.34<br>16kHz: 48.00 ± 5.61<br>24kHz: 57.00 ± 6.24<br>32kHz: 68.00 ± 4.90 | Click: 32.50 ± 1.12<br>8kHz: 48.33 ± 3.33<br>16kHz: 39.17 ± 2.01<br>24kHz: 46.67 ± 3.57<br>32kHz: 59.17 ± 4.36 |
| WT HL | Click: 56.90 ± 2.57<br>8kHz: 69.52 ± 2.39<br>16kHz: 55.95 ± 2.75<br>24kHz: 66.67 ± 3.19<br>32kHz: 76.90 ± 3.48 | Click: 54.69 ± 2.64<br>8kHz: 68.12 ± 2.95<br>16kHz: 54.38 ± 2.37<br>24kHz: 61.56 ± 3.59<br>32kHz: 70.00 ± 3.93 | Click: 54.00 ± 4.23<br>8kHz: 71.33 ± 3.98<br>16kHz: 58.67 ± 3.98<br>24kHz: 64.67 ± 4.13<br>32kHz: 78.33 ± 2.66 |
| WT NH | Click: 39.55 ± 2.90<br>8kHz: 53.18 ± 2.63<br>16kHz: 49.09 ± 3.08<br>24kHz: 52.27 ± 2.81<br>32kHz: 58.64 ± 3.02 | Click: 29.17 ± 1.54<br>8kHz: 50.83 ± 3.00<br>16kHz: 40.00 ± 3.16<br>24kHz: 45.00 ± 3.65<br>32kHz: 52.50 ± 4.61 | Click: 35.00 ± 2.24<br>8kHz: 49.00 ± 4.00<br>16kHz: 45.00 ± 4.74<br>24kHz: 55.00 ± 3.16<br>32kHz: 64.00 ± 3.67 |

## References

1. Javitt DC, Sweet RA (2015): Auditory dysfunction in schizophrenia: integrating clinical and basic features. Nat Rev Neurosci 16: 535–550.

2. Viertiö S, Perälä J, Saarni S, Koskinen S, Suvisaari J (2014): Hearing loss in persons with psychotic disorder–findings from a population-based survey. Schizophrenia Research 159: 309–311.

3. Saperstein AM, Meyler S, Golub JS, Medalia A (2023): Correlates of hearing loss among adults with schizophrenia. Schizophrenia Research 257: 1–4.

4. Linszen MMJ, Brouwer RM, Heringa SM, Sommer IE (2016): Increased risk of psychosis in patients with hearing impairment: Review and meta-analyses. Neurosci Biobehav Rev 62: 1–20.

5. Tallal P, Piercy M (1973): Defects of Non-Verbal Auditory Perception in Children with Developmental Aphasia. Nature 241: 468–469.

6. Merzenich MM, Jenkins WM, Johnston P, Schreiner C, Miller SL, Tallal P (1996): Temporal Processing Deficits of Language-Learning Impaired Children Ameliorated by Training. Science 271: 77–81.

7. Mauk MD, Buonomano DV (2004): THE NEURAL BASIS OF TEMPORAL PROCESSING Annual Review of Neuroscience 27: 307–340.

8. Iliadou VV, Apalla K, Kaprinis S, Nimatoudis I, Kaprinis G, Iacovides A (2013): Is central auditory processing disorder present in psychosis? Am J Audiol 22: 201–208.

9. Moschopoulos N, Nimatoudis I, Kaprinis S, Sidiras C, Iliadou V (2020): Auditory processing disorder may be present in schizophrenia and it is highly correlated with formal thought disorder. Psychiatry Research 291: 113222.

10. Fitzgibbons PJ, Wightman FL (1982): Gap detection in normal and hearing-impaired listeners. J Acoust Soc Am 72: 761–765.

11. Boot E, Óskarsdóttir S, Loo JCY, Crowley TB, Orchanian-Cheff A, Andrade DM, et al. (2023): Updated clinical practice recommendations for managing adults with 22q11.2 deletion syndrome. Genetics in Medicine 25: 100344.

12. Bassett AS, Chow EWC (2008): Schizophrenia and 22q11.2 deletion syndrome. Curr Psychiatry Rep 10: 148–157.

13. Verheij E, Derks LSM, Stegeman I, Thomeer HGXM (2017): Prevalence of hearing loss and clinical otologic manifestations in patients with 22q11.2 deletion syndrome: A literature review. Clinical Otolaryngology 42: 1319–1328.

14. Fuchs JC, Zinnamon FA, Taylor RR, Ivins S, Scambler PJ, Forge A, et al. (2013): Hearing loss in a mouse model of 22q11.2 deletion syndrome. ((Y. Herault, editor)). PLoS ONE 8: e80104.

15. Lu C, Linden JF (2025): Auditory evoked-potential abnormalities in a mouse model of 22q11.2 Deletion Syndrome and their interactions with hearing impairment. Transl Psychiatry 15: 1–13.

16. Kopp-Scheinpflug C, Sinclair JL, Linden JF (2018): When sound stops: Offset responses in the auditory system. Trends in Neurosciences 41: 712–728.

17. Zinnamon FA, Harrison FG, Wenas SS, Liu Q, Wang KH, Linden JF (2023): Increased Central Auditory Gain and Decreased Parvalbumin-Positive Cortical Interneuron Density in the *Df1/+* Mouse Model of Schizophrenia Correlate With Hearing Impairment. Biol Psychiatry Glob Open Sci 3: 386–397.

18. Fuchs JC, Linden JF, Baldini A, Tucker AS (2015): A defect in early myogenesis causes otitis media in two mouse models of 22q11.2 deletion syndrome. Human Molecular Genetics 24: 1869–1882.

19. Tucci DL, Cant NB, Durham D (1999): Conductive hearing loss results in a decrease in central auditory system activity in the young gerbil. The Laryngoscope 109: 1359–1371.

20. Mai J, Gargiullo R, Zheng M, Esho V, Hussein OE, Pollay E, et al. (2024): Sound-seeking before and after hearing loss in mice. Sci Rep 14: 19181.

21. Green DB, Mattingly MM, Ye Y, Gay JD, Rosen MJ (2017): Brief Stimulus Exposure Fully Remediates Temporal Processing Deficits Induced by Early Hearing Loss. J Neurosci 37: 7759–7771.

22. Keller CH, Kaylegian K, Wehr M (2018): Gap encoding by parvalbumin-expressing interneurons in auditory cortex. Journal of Neurophysiology 120: 105–114.

23. Radziwon KE, Stolzberg DJ, Urban ME, Bowler RA, Salvi RJ (2015): Salicylate-Induced Hearing Loss and Gap Detection Deficits in Rats. Front Neurol 6. 10.3389/fneur.2015.00031

24. Zinkstok JR, Boot E, Bassett AS, Hiroi N, Butcher NJ, Vingerhoets C, et al. (2019): Neurobiological perspective of 22q11.2 deletion syndrome. The Lancet Psychiatry 6: 951–960.

25. Nehme R, Pietiläinen O, Artomov M, Tegtmeyer M, Valakh V, Lehtonen L, et al. (2022): The 22q11.2 region regulates presynaptic gene-products linked to schizophrenia. Nat Commun 13: 3690.

26. Khan TA, Revah O, Gordon A, Yoon S-J, Krawisz AK, Goold C, et al. (2020): Neuronal defects in a human cellular model of 22q11.2 deletion syndrome. Nat Med 26: 1888–1898.

27. Chun S, Westmoreland JJ, Bayazitov IT, Eddins D, Pani AK, Smeyne RJ, et al. (2014): Specific disruption of thalamic inputs to the auditory cortex in schizophrenia models. Science 344: 1178–1182.

## References

1. Lindsay EA, Botta A, Jurecic V, Carattini-Rivera S, Cheah Y-C, Rosenblatt HM, et al. (1999): Congenital heart disease in mice deficient for the DiGeorge syndrome region. Nature 401: 379–383.

2. Fuchs JC, Linden JF, Baldini A, Tucker AS (2015): A defect in early myogenesis causes otitis media in two mouse models of 22q11.2 deletion syndrome. Human Molecular Genetics 24: 1869–1882.

3. Fuchs JC, Zinnamon FA, Taylor RR, Ivins S, Scambler PJ, Forge A, et al. (2013): Hearing loss in a mouse model of 22q11.2 deletion syndrome. ((Y. Herault, editor)). PLoS ONE 8: e80104.

4. Tucci DL, Cant NB, Durham D (1999): Conductive hearing loss results in a decrease in central auditory system activity in the young gerbil. The Laryngoscope 109: 1359–1371.

5. Mai J, Gargiullo R, Zheng M, Esho V, Hussein OE, Pollay E, et al. (2024): Sound-seeking before and after hearing loss in mice. Sci Rep 14: 19181.

6. Lu C, Linden JF (2025): Auditory evoked-potential abnormalities in a mouse model of 22q11.2 Deletion Syndrome and their interactions with hearing impairment. Transl Psychiatry 15: 1–13.

7. Buccino AP, Hurwitz CL, Garcia S, Magland J, Siegle JH, Hurwitz R, Hennig MH (2020): SpikeInterface, a unified framework for spike sorting. eLife 9: e61834.

8. Pachitariu M, Steinmetz N, Kadir S, Carandini M, Kenneth D. H (2016): *Kilosort: Realtime Spike-Sorting for Extracellular Electrophysiology with Hundreds of Channels*. Neuroscience. 10.1101/061481

9. Resnik J, Polley DB (2017): Fast-spiking GABA circuit dynamics in the auditory cortex predict recovery of sensory processing following peripheral nerve damage. eLife 6: e21452.

10. Li L-y., Xiong XR, Ibrahim LA, Yuan W, Tao HW, Zhang Li (2015): Differential receptive field properties of parvalbumin and somatostatin inhibitory neurons in mouse auditory cortex. Cereb Cortex 25: 1782–1791.

11. Sun SH, Almasi A, Yunzab M, Zehra S, Hicks DG, Kameneva T, et al. (2021): Analysis of extracellular spike waveforms and associated receptive fields of neurons in cat primary visual cortex. The Journal of Physiology 599: 2211–2238.

